# A Heart Disease-Associated TSPO Variant Alters Transmembrane Helix Dynamics

**DOI:** 10.64898/2026.04.10.717289

**Authors:** Aleksandra Kusova, Gwladys Rivière, Karin Giller, Marion Laudette, Jan Borén, Stefan Becker, Markus Zweckstetter

## Abstract

The 18-kDa translocator protein TSPO is an outer mitochondrial membrane protein involved in cholesterol transport, stress response, and cellular metabolism. Although its five-helix architecture is conserved across species, the structural and dynamic features of human TSPO remain unresolved. Using solution NMR spectroscopy, we characterize the conformational dynamics of human TSPO in complex with a third-generation diagnostic ligand. We identify a dynamic N-terminal segment of the first transmembrane helix that does not form a stably populated helix but instead defines a flexible boundary between the cytosolic region and the transmembrane core. The common disease-associated A14V variant reduces this conformational heterogeneity by introducing short-range contacts and redistributing backbone dynamics across the protein. These changes preserve the overall fold while locally stabilizing the N-terminal transmembrane helix toward the VDAC interaction interface. Our findings reveal a human-specific dynamic architecture of TSPO and link variant-induced stabilization to modulation of transmembrane helix dynamics.

## Introduction

The 18-kDa translocator protein (TSPO) is an outer mitochondrial membrane protein (Fig. 1A) initially described as a peripheral benzodiazepine receptor. It participates in a variety of essential processes, including cholesterol transport, steroidogenesis, immune signaling, mitochondrial permeability transition, and apoptosis (*1–4*). TSPO binds a wide range of ligands – from synthetic drugs like isoquinoline carboxamides and benzodiazepines to endogenous molecules such as porphyrins and the diazepam binding inhibitor (*5–8*).

**Fig. 1.**
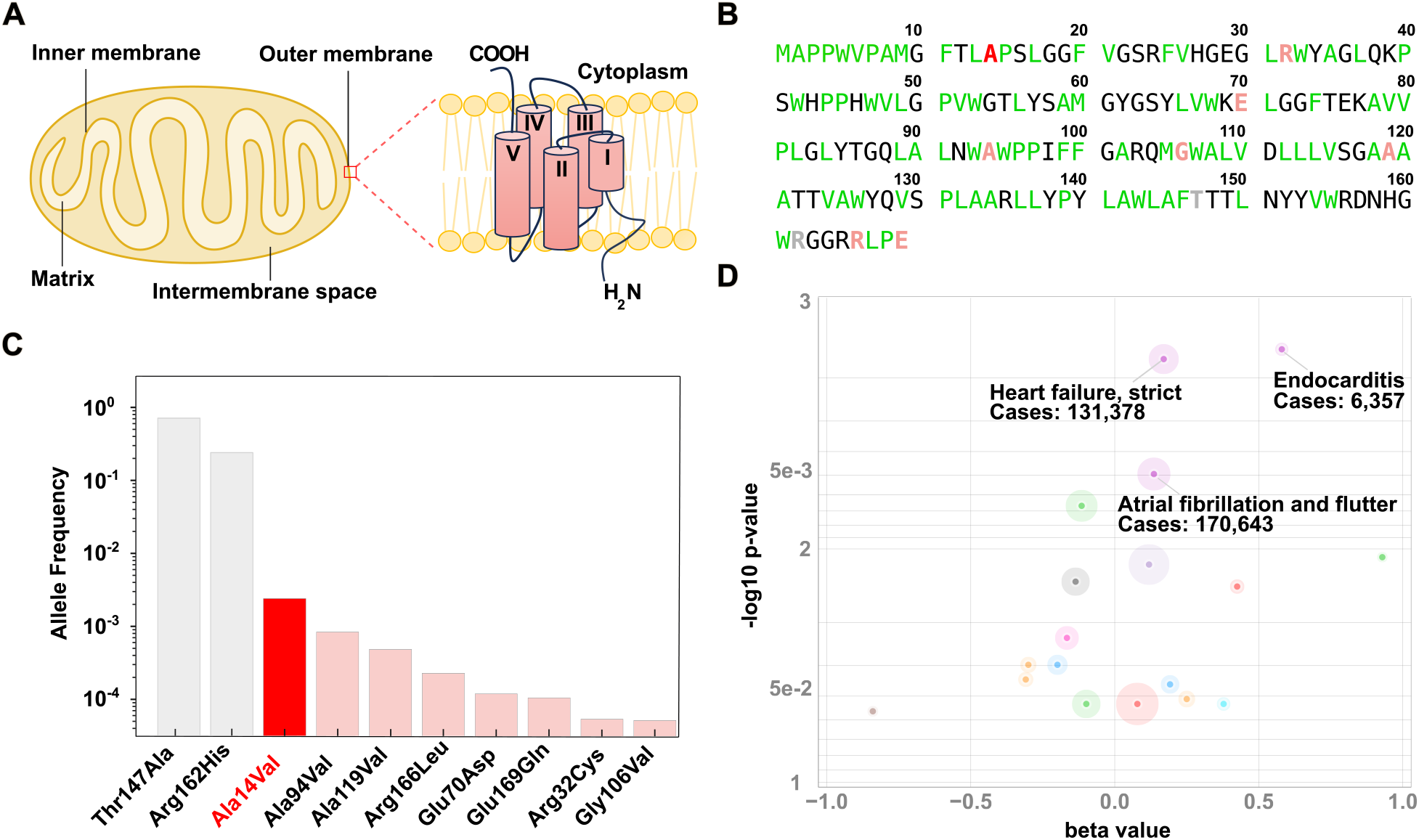
Topology, genetic variability, and clinical associations of human TSPO. (**A**) Schematic topology of hTSPO (TM1-TM5) in the outer mitochondrial membrane. (**B**) Amino acid sequence of hTSPO with hydrophobic residues (green) and frequent variant positions color-coded according to minor allele frequency. (**C**) Minor allele frequencies of the ten most common TSPO variants (gnomAD). Common alleles Thr147Ala and Arg162His are shown in gray; rare missense variants in pink with A14V (red) being the most frequent. (**D**) Top disease associations for the A14V variant based on meta-analysis from FinnGen and UK Biobank.

In the central nervous system, TSPO is expressed in glial, endothelial, and neuronal cells (*4*), and is upregulated in activated microglia and astrocytes, making it a widely used – though not cell-type–specific – marker of neuroinflammation in clinical PET imaging and preclinical models (*9*, *10*). Beyond the central nervous system, TSPO is abundantly expressed in peripheral tissues with high metabolic demand, including the heart, where it has been implicated in mitochondrial function, cholesterol handling, oxidative stress responses, and cardiomyocyte survival (*11*, *12*). Dysregulation of these processes is associated with heart failure, arrhythmogenesis, and ischemic injury, highlighting potential relevance of TSPO function in cardiovascular physiology (*13*, *14*).

Despite its broad physiological relevance, the molecular mechanisms by which TSPO exerts its functions remain incompletely understood. TSPO’s conformational variability and oligomerization have been proposed to contribute to functional regulation, yet the structural basis of these processes is poorly defined. While the overall fold – five transmembrane helices arranged in a compact bundle – appears conserved across species (Fig. 1A, Fig. S1A) (*15–18*), detailed studies have revealed pronounced differences in flexibility, ligand binding, and oligomeric state across species (*19*, *20*). These observations raise the possibility that dynamic features, rather than static architecture alone, may underlie species-specific aspects of TSPO function.

This structural variability is particularly relevant for human TSPO (hTSPO), which harbors several naturally occurring missense variants (Fig. 1B). Among these, the A14V variant is the most frequent missense polymorphism according to gnomAD (Fig. 1C) (*21*). Large-scale association studies, including FinnGen, link A14V to increased risk of cardiovascular conditions such as heart failure and atrial fibrillation (Fig. 1D), with cardiovascular and cerebrovascular phenotypes emerging as the strongest individual signals (Fig. S1B) (*21*). However, how this common variant influences the structural and dynamic properties of hTSPO remains unknown (*20*).

In this study, we present a structural and dynamic characterization of hTSPO using solution NMR spectroscopy. We identify a highly flexible N-terminal segment of the first transmembrane helix that contrasts with the stable helical architecture observed in other species. We further show that the common A14V polymorphism, located at the boundary between this dynamic segment and the transmembrane core, reduces conformational heterogeneity without perturbing the global fold, revealing how a frequent human variant reshapes the dynamic landscape of TSPO.

## Results

### Flexible TM1-N segment in hTSPO

To define the structural basis of human TSPO, we recombinantly prepared hTSPO and reconstituted it together with the diagnostic PET ligand GE-180 into either DPC micelles or (DMPC/DHPC, *q* = 0.5) bicelles (Fig. S2A). The [^1^H,^15^N]-TROSY spectra of the hTSPO/GE-180 complex in both membrane-mimicking environments exhibit well-dispersed cross peaks in contrast to ligand-free hTSPO (Fig. 2A and Fig. S2B). Chemical-shift perturbations between the two conditions were generally small and did not cluster in specific regions, indicating that the protein adopts a well-folded state in both environments (Fig. S2C). Observed deviations were mostly restricted to residues at the detergent-water interface, pointing to environmental adjustments rather than genuine structural rearrangements.

**Fig. 2.**
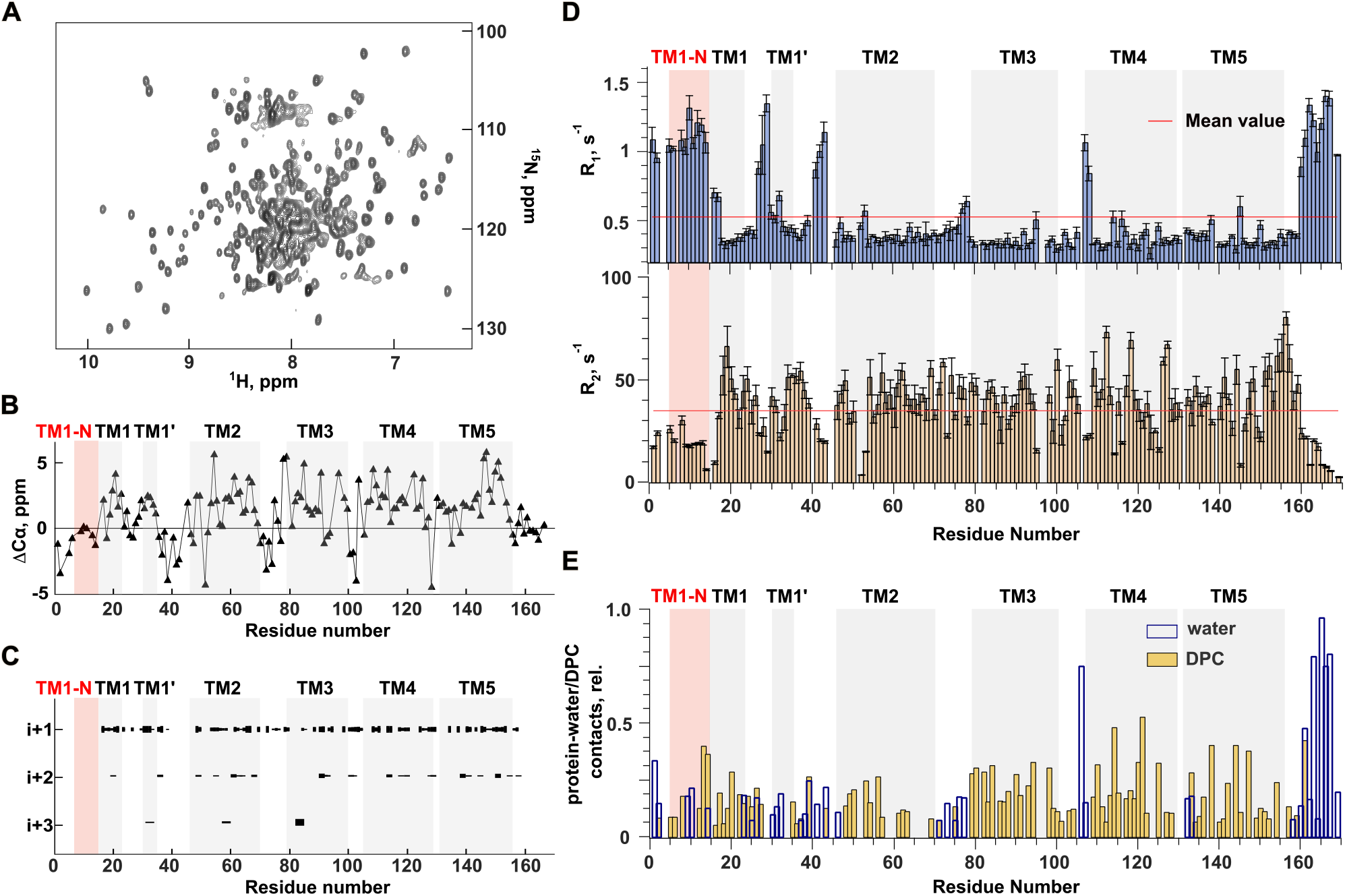
Structural dynamics in human TSPO. (**A**) 2D [^1^H,^15^N]-TROSY spectrum of hTSPO in complex with the GE-180 in DPC micelles. (**B**) Secondary chemical shifts (ΔCα) of hTSPO. (**C**) Sequential amide-amide contacts (i→i+1 to i→i+3). The absence of contacts in TM1-N indicates local disorder. (**D**) Residue-specific ^15^N longitudinal (R_1_) and transverse (R_2_) relaxation rates in hTSPO. (**E**) Residue-specific water and DPC contact profiles derived from cross-peak intensities in the 3D ^15^N-edited NOESY-TROSY spectrum.

Using multidimensional NMR experiments, we assigned 95% of the backbone resonances of 161-residue hTSPO in complex with GE-180 in DPC micelles (Fig. 2B). Secondary chemical shifts together with sequential i→i+1; i+2 NOE cross peaks unambiguously defined the position of the four transmembrane helices TM2 to TM5. In contrast, residues from V6 up to A14 that form the N-terminal part of TSPO’s first transmembrane region (TM1-N) lacked helix-specific NOE contacts (Fig. 2C), suggesting that TM1-N does not adopt a persistently populated helical conformation on the NMR time scale.

To confirm this distinct feature of TM1, we next examined backbone dynamics. We measured ^15^N longitudinal (R_1_) and transverse (R_2_) relaxation rates using TROSY-optimized pulse sequences that are designed for large proteins and membrane proteins (*22*). The transmembrane domains show the characteristic pattern of higher R_2_ and lower R_1_ values, indicative of restricted motions, whereas TM1-N displays reduced R_2_ and elevated R_1_ values comparable to the flexible N- and C-termini (Fig. 2D). This behavior demonstrates that TM1-N exhibits enhanced backbone mobility consistent with the absence of a stably populated secondary structure.

Further support for a more disordered potentially dynamic state of the N-terminal half of TM1 was provided by intermolecular NOEs between backbone amide groups and water (Fig. S3). hTSPO’s termini and interhelical loops displayed pronounced NOE-detected water contacts in both DPC micelles and DMPC/DHPC bicelles (Fig. S2D-E). In contrast, contacts between backbone amide groups and DPC aliphatic protons, but not water protons, were observed for residues in hTSPO’s transmembrane helices in agreement with their location in a hydrophobic environment. However, residues in TM1-N showed both water and DPC NOEs (Fig. 2E), indicating that TM1-N can engage in detergent contacts while maintaining amide–water NOE interactions (Fig. S4).

The single-residue analysis demonstrates that hTSPO retains the canonical five-helix bundle, while incorporating a locally disordered 9-residue N-terminal segment of TM1 that exhibits amide–water NOE contacts. In contrast, the corresponding TM1-N segment in mouse TSPO (mTSPO) displays a well-formed α-helix (Fig. S1C). This suggests that the observed conformational plasticity of TM1-N may represent a species-specific structural feature with potential regulatory implications.

### hTSPO displays multiple conformations in TM1

To investigate the structural basis of the dynamic heterogeneity in hTSPO, we first analyzed the product R_1_R_2_ for residues with high hetNOE values, which report on restricted backbone motions (Fig. S5). This approach revealed sites with elevated R_1_R_2_ values not only in loops and termini but also within TM1-N and TM4 structural regions, consistent with contributions from conformational exchange on the μs-ms timescale (Fig. 3A).

**Fig. 3.**
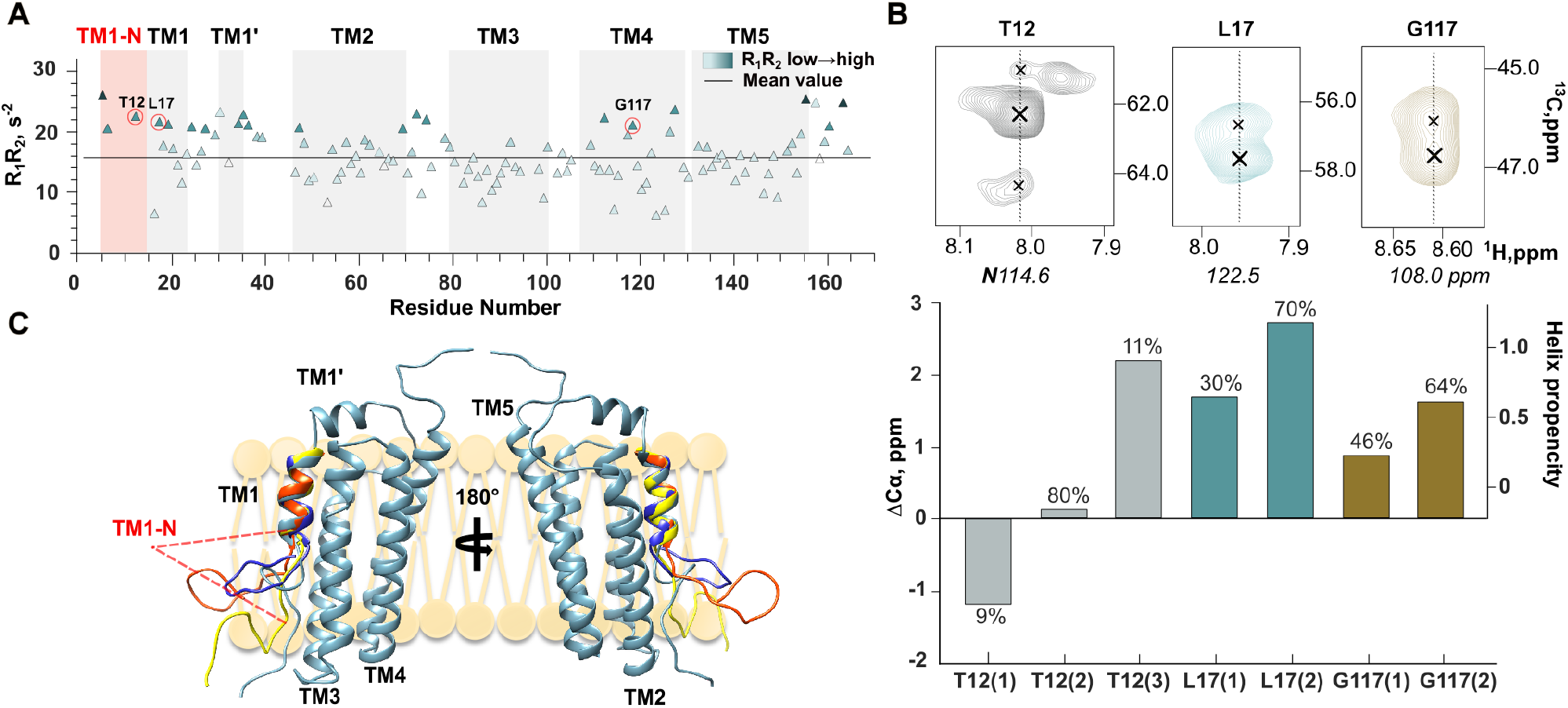
Conformational heterogeneity of TM1-N in hTSPO. (**A**) R_1_R_2_ values along the hTSPO sequence, color-coded from low to high. Red circles indicate residues for which both major and minor Cα resonances were detected. (**B**) Major and minor Cα resonances of T12, L17, and G117 (top). Secondary Cα chemical shifts of major and minor states with relative populations estimated from integral peak intensities (bottom). (**C**) Structural model of hTSPO showing the position of TM1-N. The TM1 ensemble includes four representative conformers of the major Cα chemical-shift state, embedded within the full-length hTSPO structure.

Supporting this interpretation, additional minor resonances were detected in the Cα planes of 3D HNCA spectra for residues in both TM1-N and TM4 (Fig. 3B top), indicating the presence of alternative backbone conformations. To estimate the structural nature of the observed minor states, we calculated ΔCα-based helix propensities. The secondary conformers at L17 and G117 retained helical character, consistent with the dominant state. In contrast, one of the minor resonances shifted into the α-helical regime for T12 in TM1-N (Fig. 3B bottom) corresponding to a low-populated (∼11%) minor state, based on relative peak intensities.

To visualize the structural consequences of this heterogeneity, we generated an ensemble of TM1 conformations based on experimental chemical shifts and TALOS+ predictions and combined it with the 3D structure of hTSPO predicted by AlphaFold3. In the resulting model, which is intended as a qualitative visualization of experimentally observed conformational heterogeneity, the helical part of TM1 is well-defined while the TM1-N segment samples multiple conformations oriented toward the membrane surface (Fig. 3C).

### A14V mutation decreases conformational heterogeneity in TM1

The structural heterogeneity in the N-terminal half of TM1 is specific to hTSPO, while the same segment in mTSPO folds into a stable α-helix (Fig. S1C). Comparison of the amino acid sequences of this segment between hTSPO and mTSPO reveals the amino acid substitutions M9V, F11L and A14V (Fig. 4A). The A14V substitution is located at the edge of the helical structure in the TM1 region of hTSPO and is the only one of the three substitutions that corresponds to a clinically relevant variant (Fig. 1C,D).

**Fig. 4.**
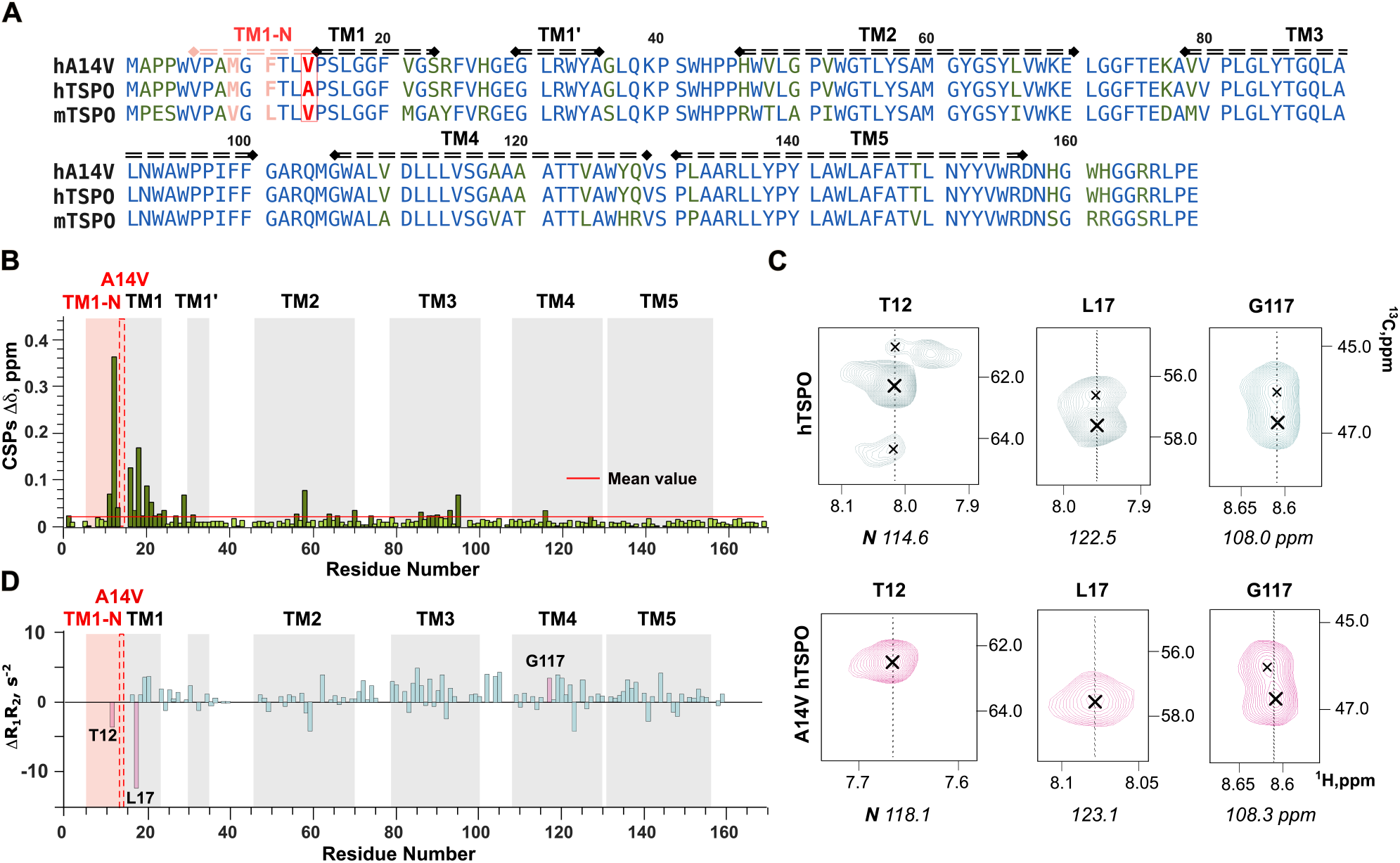
A14V eliminates conformational heterogeneity of TM1. (**A**) Sequence alignment of hTSPO, A14V hTSPO, and mTSPO. Conserved residues are shown in blue, species-specific substitutions in green, the TM1-N segment in rose, and the A14V substitution in red. (**B**) Chemical-shift perturbations (CSPs) between wild-type and A14V hTSPO. The red dashed line indicates the position of the A14V substitution. (**C**) Cα chemical shifts for selected residues (T12, L17, G117) in wild-type (top) and A14V hTSPO (bottom). (**D**) ΔR_1_R_2_ between A14V and wild-type hTSPO along the sequence. Residues included in this analysis were filtered based on ^15^N-hetNOE data (Fig. S7).

We therefore examined how the A14V substitution influences the structure of hTSPO. Superposition of the [^1^H,^15^N]-TROSY spectra of wild-type and A14V hTSPO (Fig. S6A) exhibits similar overall dispersion, confirming that the global fold is preserved, but several resonances near the mutation site show pronounced chemical-shift changes (Fig. S6B). Mapping these perturbations onto the sequence (Fig. 4B) demonstrates that the strongest effects are localized to TM1-N and TM1, while smaller but consistent shifts appear in TM2 and TM3 residues spatially adjacent to the mutation (Fig. S6C). This pattern indicates that A14V modifies the local environment of TM1 and subtly influences the packing of neighboring regions.

The analysis of secondary chemical shifts (ΔCα) revealed that in the TM1 region, where the mutation is located, the minor conformational states detected in the wild-type protein disappeared, leaving a single dominant conformation for residues T12 and L17 (Fig. 4C). In contrast, the conformational heterogeneity of G117 within TM4 persisted, indicating that the effect of the mutation is local. Consistently, 1′R_1_R_2_ analysis revealed negative values for T12 and L17 in A14V hTSPO, consistent with a reduced contribution from conformational exchange in TM1, while other regions remained largely unaffected (Fig. 4D).

### A14V stabilizes TM1 and affects connected regions

To further characterize the structural consequences of the A14V substitution, we compared the sequential amide-amide contacts detected for wild-type and A14V hTSPO from the 3D ^15^N-edited NOESY-TROSY spectra (Fig. 5A). Overall, the contact pattern was largely similar between the two proteins, indicating that the global topology remained preserved. However, in the TM1-N, where the mutation is located, A14V hTSPO exhibited additional i→i+1 contacts that were absent in the wild-type hTSPO. The appearance of these short-range interactions indicates reduced conformational heterogeneity and increased structural definition of TM1-N in A14V hTSPO.

**Fig. 5.**
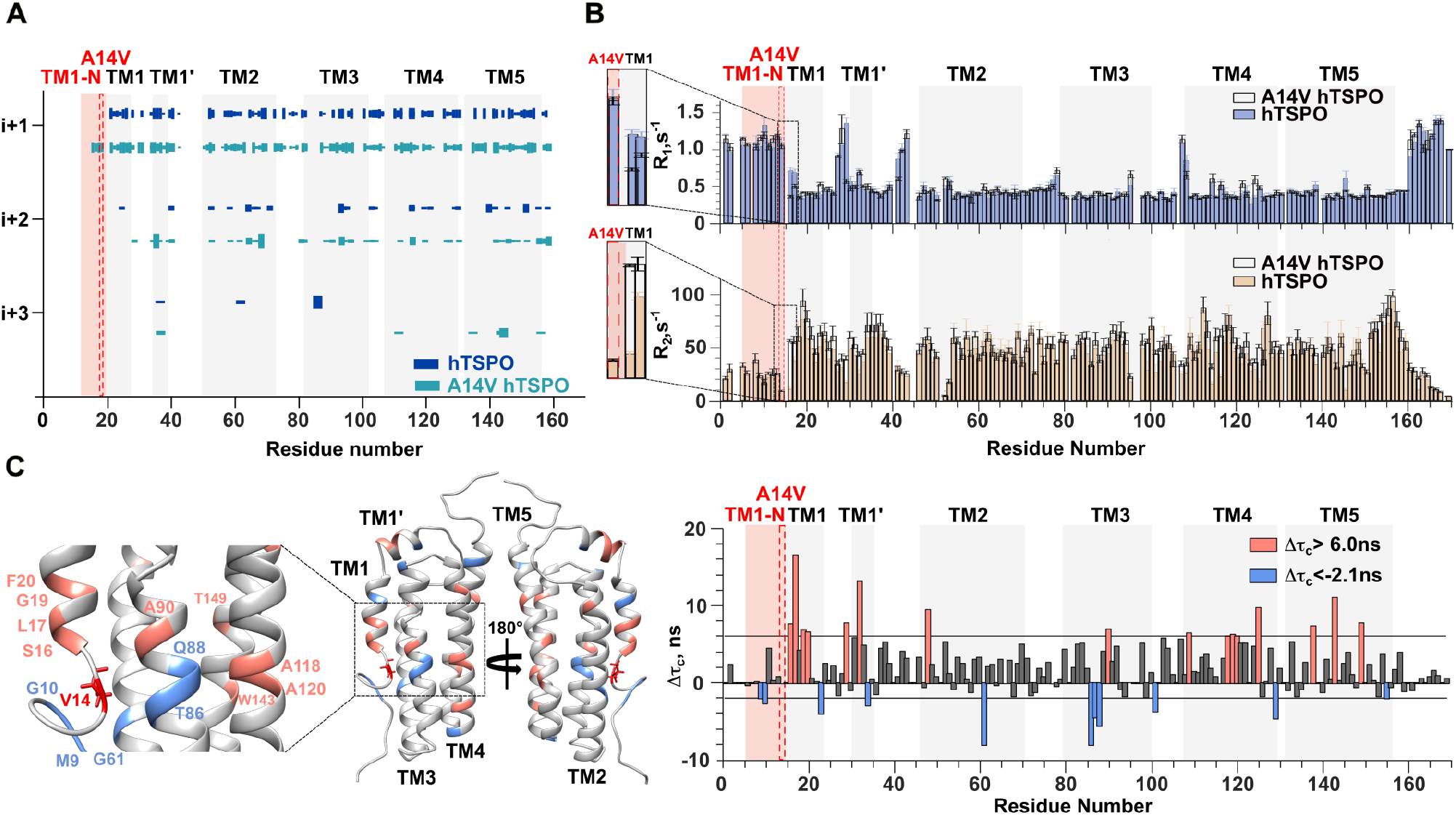
A14V is associated with reduced TM1 conformational heterogeneity and altered backbone dynamics. (**A**) Sequential amide–amide contacts (i→i+1 to i→i+3) observed for wild-type and A14V hTSPO in complex with GE-180 in DPC micelles. TM1-N lacks detectable contacts in wild-type hTSPO, whereas i→i+1 contacts are observed in the A14V variant. (**B**) Residue-specific ^15^N longitudinal (R_1_) and transverse (R_2_) relaxation rates of A14V hTSPO in comparison with the wild-type protein. (**C**) Residue-specific differences in backbone rotational correlation times (Δτ_c_ = τ_c_(A14V) – τ_c_(wild-type)) mapped onto the structure (left) and shown as a bar plot (right); salmon indicates values above 6.0 ns and light blue values below –2.1 ns.

Analysis of the ^15^N relaxation rates revealed reduced R_1_ and increased R_2_ values for residues 16 and 17 within TM1 upon the A14V substitution (Fig. 5B). This indicates that the mutation locally stabilizes this part of TM1 while other parts of the protein remain largely unchanged. To evaluate global effects of the mutation, we additionally compared residue-specific amide–water NOE profiles between wild-type and A14V hTSPO (Fig. S8A). The highly similar profiles indicate that the micelle–water interface and overall water proximity remain largely unchanged upon mutation.

Analysis of backbone rotational correlation times (τ_c_) revealed that the A14V substitution is associated with changes in backbone dynamics across several transmembrane helices (Fig. 5C). TM1 exhibits a pronounced increase in τ_c_, reflecting local rigidification, particularly for residues S16-F20. Additional regions with elevated τ_c_ values include A90 (TM3), A118/A120 (TM4), and W143/T149 (TM5), indicating mutation-induced slowing of backbone motions across multiple helices. In contrast, G61 and T86-Q88 (TM2-TM3) show decreased τ_c_ values, consistent with locally enhanced flexibility. When mapped onto the structure, many of these changes cluster at a similar axial height within the transmembrane bundle, suggesting a lateral redistribution of backbone dynamics triggered by the A14V mutation. On average, τ_c_ increased from 24.1 ns in wild-type hTSPO to 25.7 ns in A14V hTSPO (Fig. S8B), consistent with a modest reduction in overall backbone mobility. Together, these results show that A14V promotes local ordering through formation of additional short-range contacts in TM1-N and reduced backbone flexibility in the adjacent TM1 segment, while causing a redistribution of dynamics across the transmembrane bundle.

## Discussion

Our study reveals an unexpected and human-specific feature in the architecture of TSPO: the N-terminal segment of the first transmembrane helix (TM1-N) does not form a persistently populated α-helix but instead exhibits pronounced conformational heterogeneity at the membrane interface. This behavior contrasts with TSPO homologs from other species, including mouse TSPO, where the corresponding region adopts a stable transmembrane helix (*16–19, 23*). Despite this local deviation, the overall five-helix topology of TSPO is preserved in both detergent micelles and bicelles, underscoring that TM1-N represents a localized structural specialization rather than a global rearrangement.

Multiple independent NMR observables consistently define TM1-N as a dynamic membrane-interfacial region. Secondary-structure propensities derived from chemical shifts confirm stable helices for TM2-TM5 while revealing a clear discontinuity at TM1-N (Fig. 6A). Relaxation data further distinguish this region by elevated R_1_ and reduced R_2_ values relative to the transmembrane core, indicative of enhanced backbone mobility (Fig. 2D). Intermolecular NOEs demonstrate that TM1-N simultaneously contacts both detergent and water (Fig. 2E), placing it at the membrane interface rather than fully embedded within the hydrophobic core. Together, these data define TM1-N as a partially inserted, conformationally heterogeneous segment that bridges the disordered cytosolic tail and the transmembrane bundle. Notably, this experimentally defined architecture diverges from AlphaFold3 predictions, which favor a continuous helix in this region, highlighting a limitation of static structure prediction in capturing membrane-interfacial dynamics (Fig. 6A).

**Fig. 6.**
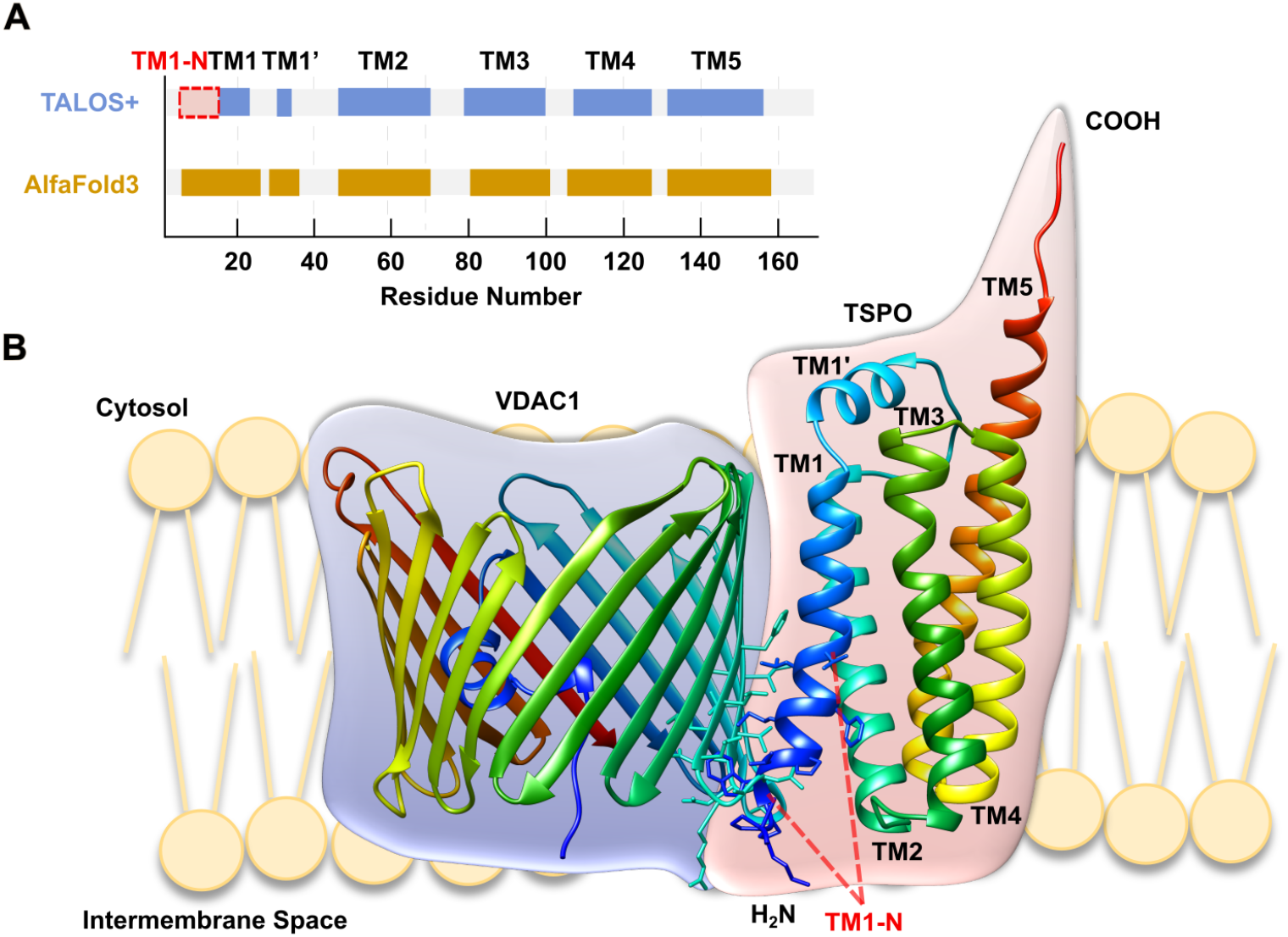
TM1-N defines a dynamic membrane-interfacial region in hTSPO. (**A**) TALOS+ secondary-structure propensities derived from NMR chemical shifts and AlphaFold3 predictions. (**B**) AlphaFold3-predicted model of the hTSPO–VDAC complex.

We further show that the common human TSPO variant A14V selectively modulates this dynamic landscape. Rather than inducing a new fold, A14V suppresses low-populated conformational states within TM1-N and shifts the equilibrium toward a more structurally defined ensemble. This population shift is reflected by the disappearance of minor backbone resonances (Fig. 4C), the emergence of additional short-range sequential contacts (Fig. 5A), and reduced exchange contributions for residues proximal to the mutation site (Fig. 5C). Importantly, these effects are local: the overall fold, amide–water NOE profile, and membrane topology of TSPO remain unchanged (Figs. 4B and S8), and conformational heterogeneity in distant regions such as TM4 is preserved (Fig. 4D). Thus, A14V acts as a local modulator of conformational heterogeneity rather than a global stabilizer.

Beyond TM1, cross-correlated relaxation analysis reveals that A14V is associated with altered backbone dynamics across multiple transmembrane helices. Regions exhibiting slower backbone dynamics cluster predominantly within the interior of the transmembrane bundle, whereas select segments display increased flexibility (Fig. S9). These changes do not imply rigidification of the protein as a whole but instead indicate a redistribution of backbone dynamics that preserves the global architecture. Such long-range dynamic coupling is consistent with subtle repacking effects within tightly packed membrane proteins, where local stabilization can influence motional properties of neighboring helices without altering topology.

The location and behavior of TM1-N suggest a broader mechanistic role as a membrane-interfacial boundary helix. Partially inserted or re-entrant helices are increasingly recognized as dynamic elements that facilitate folding, assembly, and regulation of membrane proteins (*24, 25*). In this context, TM1-N may act as a flexible interface during early stages of transmembrane bundle organization, allowing adaptive packing of the adjacent helical region. Stabilization of this boundary by A14V could reduce conformational sampling at the membrane interface, thereby fine-tuning local helix dynamics. Such effects are well positioned to influence processes that depend on precise transmembrane packing, including ligand accommodation near TM1 and the maturation of TSPO within the mitochondrial outer membrane.

TSPO functions within multiprotein assemblies on the outer mitochondrial membrane, most prominently in association with the voltage-dependent anion channel (VDAC) (*26–28*). Although our data do not directly probe TSPO-VDAC interactions, AlphaFold3-based models place TM1-N near the proposed interaction interface, providing spatial context for the observed dynamics (Fig. 6B). In this framework, mutation-induced stabilization of TM1-N may influence local packing at protein-protein interfaces and thereby modulate interaction propensities within mitochondrial complexes. In cardiomyocytes, where mitochondrial activity is tightly coupled to energy demand, TSPO biogenesis and maturation are further regulated by chaperone systems such as GRP78 (*13*). Local stabilization of TM1-N at the membrane boundary could, in principle, alter the balance between chaperone-associated intermediates and mature assemblies. While these possibilities remain to be tested experimentally, our structural data provide a plausible mechanistic framework linking human-specific TSPO dynamics to modulation of interaction-competent conformational states.

In summary, this work establishes TM1-N as a dynamic, human-specific boundary element in TSPO and identifies A14V as a population-shifting variant that selectively reduces conformational heterogeneity without perturbing global structure. By defining how a common disease-associated variant reshapes the dynamic landscape of a membrane protein at single-residue resolution, our study provides a structural foundation for future investigations into TSPO assembly, interaction networks, and functional regulation in native mitochondrial membranes.

## Materials and Methods

### Molecular cloning, expression, purification, and sample preparation

Wild-type human TSPO (UniProt: P30536) and its A14V variant were cloned into a modified pET16b expression vector for bacterial expression and immobilized metal affinity chromatography. The construct encodes a common hTSPO allele carrying the frequent polymorphisms T147A and R162H. To prevent disulfide bond formation and improve protein stability, cysteine residues C19 and C153 were replaced by glycine and tyrosine, respectively. The A14V substitution was introduced by site-directed mutagenesis, and all constructs were verified by DNA sequencing.

Expression of both wild-type and A14V TSPO was performed in M9 minimal medium in Escherichia coli strain BL21(DE3) at 37 °C. After induction with 1 mM isopropyl β-D-1-thiogalactopyranoside (IPTG), the expression culture was harvested after 16 hours. The wild-type protein was uniformly [^2^H, ^13^C, ^15^N]-labeled using deuterated M9 minimal medium containing ^15^NH_4_Cl and [^13^C_6_]-glucose as the sole nitrogen and carbon sources, while the A14V mutant was produced in fully perdeuterated medium ([U-^2^H, ^13^C, ^15^N]) to enhance spectral resolution.

After harvesting, the cells were resuspended in 50 mM HEPES, pH 7.8 and 150 mM NaCl and lysed by ultrasonication. After centrifugation at 5,000 × g, the supernatant was discarded and the pellet was washed repeatedly with lysis buffer, followed by solubilization in buffer with 1% (w/v) sodium dodecyl sulfate (SDS) and 625 U Benzonase (Merck) for each liter of expression medium. After another centrifugation at 48,000 × g, the protein was loaded onto Ni-NTA resin (Qiagen), washed with lysis buffer, followed by detergent exchange to 3 % (w/v) DPC on column. The protein was eluted with lysis buffer containing 6 % DPC and 250 mM imidazole.

Micellar protein samples were buffer-exchanged at room temperature to 10 mM sodium phosphate, pH 6.0 with 2 % (w/v) DPC using Zeba spin desalting columns with 7 kDa MWCO. These micellar samples were loaded with GE-180 by adding the compound sequentially from a 140 mM stock in DMSO, not exceeding 4 % (v/v) of the sample volume, to add the compound at a final ratio of GE-180/TSPO of about 200/1 (mol/mol). After each addition of GE-180 the sample was dialyzed for at least 6 hours against 10 mM sodium phosphate, pH 6.0 using a 10 kDa MWCO Slide-A-Lyzer dialysis cassette (Pierce). Finally, the sample concentration was adjusted at room temperature to 0.45 mM using an AMICON Ultra 3 kDa MWCO concentrator.

Samples intended for bicellar reconstitution were concentrated after elution from the Ni-NTA resin using an Amicon Ultra 30 kDa MWCO concentrator to a final concentration of 9.5 mg/mL. The NaCl concentration was adjusted to 800 mM and the protein was precipitated by addition of twenty volumes of ice-cold ethanol. After incubation at -20 °C overnight the precipitate was collected by centrifugation, washed three times with ice-cold water. Finally, the protein was dissolved in 10 mM sodium phosphate, pH 6.0 supplemented with 300 mM DHPC to a final theoretical protein concentration of 1 mM. After vortexing and sonication the sample was incubated at room temperature for four days on a rotator. After removing the insoluble fraction by centrifugation, the protein sample was loaded with GE-180 according to the protocol for the micellar loading. After measuring the DHPC concentration of the sample by NMR (106 mM), it was adjusted to 300 mM and DMPC was added to the sample at a DMPC/DHPC ratio of 1/2 (mol/mol). Subsequently the sample was vortexed, sonicated, incubated at 42 °C for 15 minutes and stored for two days at room temperature to form bicelles. This sample could not be further concentrated, because spontaneous precipitation occurred at higher concentration. Therefore, the protein concentration was limited to 80–150 µM.

### Structural NMR spectroscopy

NMR samples of (^2^H, ^13^C, ^15^N)-labeled hTSPO (wild-type and A14V) in complex with GE-180 were prepared in 10 mM sodium phosphate buffer, pH 5.9, containing 0.05% (w/v) sodium azide and 10% D₂O, and reconstituted either in DPC micelles or isotropic DMPC/DHPC bicelles (*q* = 0.5). NMR spectra were recorded at 42°C on Bruker Avance III/NEO spectrometers (600, 800, 900, 950, and 1200 MHz) equipped with cryogenic TCI probes (Table S1).

For the bicellar sample, the substantially lower protein concentration limited the feasible experiments to 2D [^1^H, ^15^N]-TROSY-HSQC and 3D ^15^N-edited NOESY-TROSY spectra, both of which showed reduced overall intensity. All spectra were processed in TopSpin 3.6.3 / 4.3.0 (Bruker BioSpin) and NMRFx Analyst 11.4.1 (*29*), and analyzed using CCPNMR Analysis 3.2.2.1 (*30*). A complete list of software and versions used in this study is provided in Table S2.

Backbone resonance assignment was achieved using TROSY-based triple-resonance experiments (HNCO, HN(CA)CO, HNCA, HN(CO)CA, HNCACB). Assignments proceeded through iterative FLYA-assisted cycles (*31*): after each round, high-confidence chemical shifts were fixed as constraints, the remaining ones re-predicted, and results manually validated. In total, 16 FLYA–manual refinement cycles were performed, yielding ∼95% completeness for both wild-type and A14V hTSPO.

Intermolecular and sequential amide–amide contacts were identified from 3D ^15^N-edited NOESY-TROSY spectra (mixing time 120 ms) recorded for both wild-type and A14V hTSPO. Cross-peaks corresponding to i→i+1 to i→i+3 connectivities were manually verified and used to define local secondary-structure elements within the transmembrane helices. In addition, intermolecular NOEs to water (∼4.7 ppm) and DPC aliphatic protons (1.1–1.3 ppm) were analyzed from the same dataset to assess relative water and detergent proximity. The resulting water- and DPC-contact profiles revealed strong water NOEs for polar residues near the micelle–water interface, whereas hydrophobic residues within the transmembrane segments exhibited pronounced contacts with detergent.

### Structure calculation and AF3 modeling

Backbone chemical shifts (HN, N, C′, Cα, Cβ) were used to derive φ/ψ dihedral angles with TALOS+(*32*), and the resulting secondary-structure propensities were applied as soft restraints during structure calculation of the TM1 segment. NOE cross-peaks corresponding to short- and medium-range contacts were manually verified in TopSpin and CCPNMR, and their intensities were calibrated and classified into distance ranges: strong (1.8–3.5 Å), medium (1.8–4.5 Å), weak (1.8–5.5 Å), and very weak (1.8–6.0 Å). Hydrogen-bond restraints were included only when TALOS+ predictions were supported by characteristic amide–carbonyl NOE patterns. Structure calculations for TM1 were performed in XPLOR-NIH 3.7 (*33*) using a simulated-annealing protocol with a membrane-protein-optimized potential energy function (*34*). Structure calculations were based on the major TM1 conformation observed in the NMR spectra and yielded an ensemble consistent with partial helicity and conformational variability. To visualize TM1 within the full-length protein context, the experimental TM1 ensemble was subsequently integrated into the AF3-predicted model of hTSPO (*35*). In this composite representation, the AF3-derived transmembrane core served as a structural scaffold, while the experimentally refined TM1 ensemble reflected the conformational variability of the N-terminal region. This model was used exclusively for visualization and mapping of experimental restraints; all quantitative interpretations are based on NMR data.

Structural visualizations were prepared in UCSF Chimera 1.7 (*36*) and Chimera X (*37*). The structural model of the hTSPO–VDAC complex was generated using AlphaFold3, based on the input sequences of hTSPO and VDAC without external templates.

### Backbone dynamics

Backbone dynamics were characterized by measuring longitudinal (R_1_) and transverse (R_2_) ^15^N relaxation rates using TROSY-based pulse sequences (*22*) at 42°C on a Bruker 950 MHz spectrometer. Relaxation delays ranged from 0.20 to 3.04 s for R_1_ and from 8 to 256 ms for R_2_.

The product R_1_R_2_ was analyzed following the method of Kneller et al. (*38*), which provides a straightforward means of separating the effects of motional anisotropy and chemical exchange in relaxation data, as the contribution of the overall rotational correlation time is largely suppressed in the product term. Residues with reduced heteronuclear NOE values (below the mean: 0.62 for wild-type and 0.67 for A14V) were excluded to remove contributions from highly flexible regions. For the remaining residues, the R_1_R_2_ product was calculated individually, and mean R_1_R_2_ values of 15.9 and 16.4 s^-2^ for wild-type and A14V hTSPO, respectively, were determined over the entire sequence. Deviations from these mean values were then evaluated residue by residue. Because variations in R_1_R_2_ due to overall rotational anisotropy are minimized, elevated R_1_R_2_ values can be directly attributed to μs–ms chemical exchange. Consequently, residues exhibiting R_1_R_2_ values above the average were identified as likely participating in conformational exchange, with the magnitude of deviation correlating with the probability of exchange involvement.

Cross-correlated relaxation rates (*_gxy_*) between the ^15^N–^1^H dipolar and ^15^N chemical shift anisotropy (CSA) interactions were measured using the interleaved TROSY-HSQC pulse sequence following the implementation of Bax et al. (*39*, *40*). Two spectra were recorded with opposite ^1^H pulse phase schemes corresponding to parallel and antiparallel dipole–CSA interference pathways. The resulting difference in the apparent transverse relaxation rates of the two TROSY-HSQC components (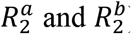) yields the cross-correlated relaxation rate:

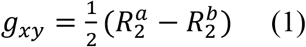

In practice, *g_xy_* is obtained from the intensity ratio of the two interleaved spectra:

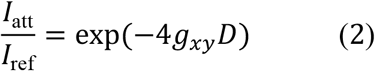

where *D* is the variable delay between the ^1^H inversion pulses during the N–H dephasing period,

*I*_att_ the attenuated signal, and *I*_ref_ the reference signal acquired under opposite dipole–CSA interference conditions.

Under the slow-tumbling regime characteristic of membrane proteins, *g_xy_* is dominated by the zero-frequency spectral density term *J*(*0*) ∝ *τ_c_*, providing a direct estimate of the rotational correlation time:

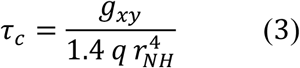

where

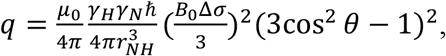

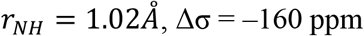, and θ ≈ 16° is the angle between the CSA and N–H bond vectors. Because the measurement is based on a difference between two transverse relaxation rates, contributions from chemical exchange and internal motions largely cancel, making this approach particularly suitable for large, anisotropically tumbling systems such as membrane proteins in detergent micelles, where model-free analysis fails due to the non-isotropic diffusion tensor and slow rotational motion. Thus, the cross-correlated relaxation method provides a direct estimate of τ_c_ that is insensitive to chemical exchange and internal motions.

## Acknowledgments

We thank Prof. Dr. Peter Bartenstein (LMU Munich) and Prof. Dr. Rupprecht (University of Regensburg) for providing the third-generation TSPO ligand GE-180. M.Z. acknowledges access to the 1.2 GHz spectrometer through the DFG Major Instrumentation Grant INST 1525/26-1 FUGG (project number 600373).

## Author contributions

Conceptualization: AK, JB, SB, MZ

Methodology: AK

Resources: KG

Investigation and visualization: AK, GR, ML

Writing and editing: AK, SB, MZ

Supervision: MZ

## Data and materials availability

All data are available in the main text or the supplementary materials.

