## Supplementary_materials for "A Heart Disease-Associated TSPO Variant Alters Transmembrane Helix Dynamics"

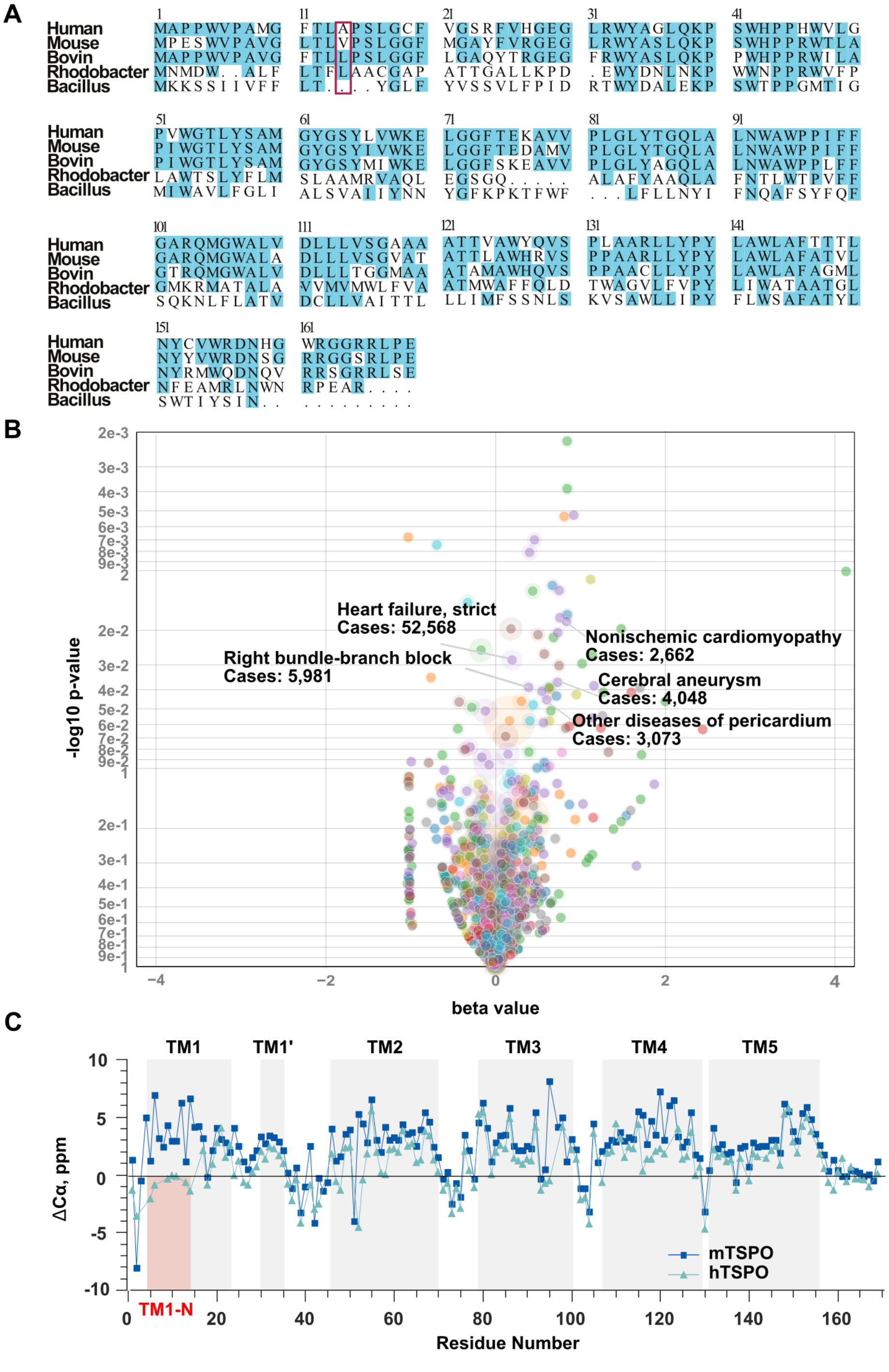


**Fig. S1.** (**A**) Multiple sequence alignment of TSPO from different species. (**B**) Phenotypic associations of the TSPO variant rs187866832 (A14V) in the FinnGen cohort (without meta-analysis**)**. Labels highlight selected cardiovascular and cerebrovascular phenotypes with relatively strong association signals in FinnGen alone. (**C**) Cα chemical-shift differences (ΔCα) between hTSPO and mTSPO. mTSPO Cα chemical shifts were obtained from BMRB entry 19608 (*16*).

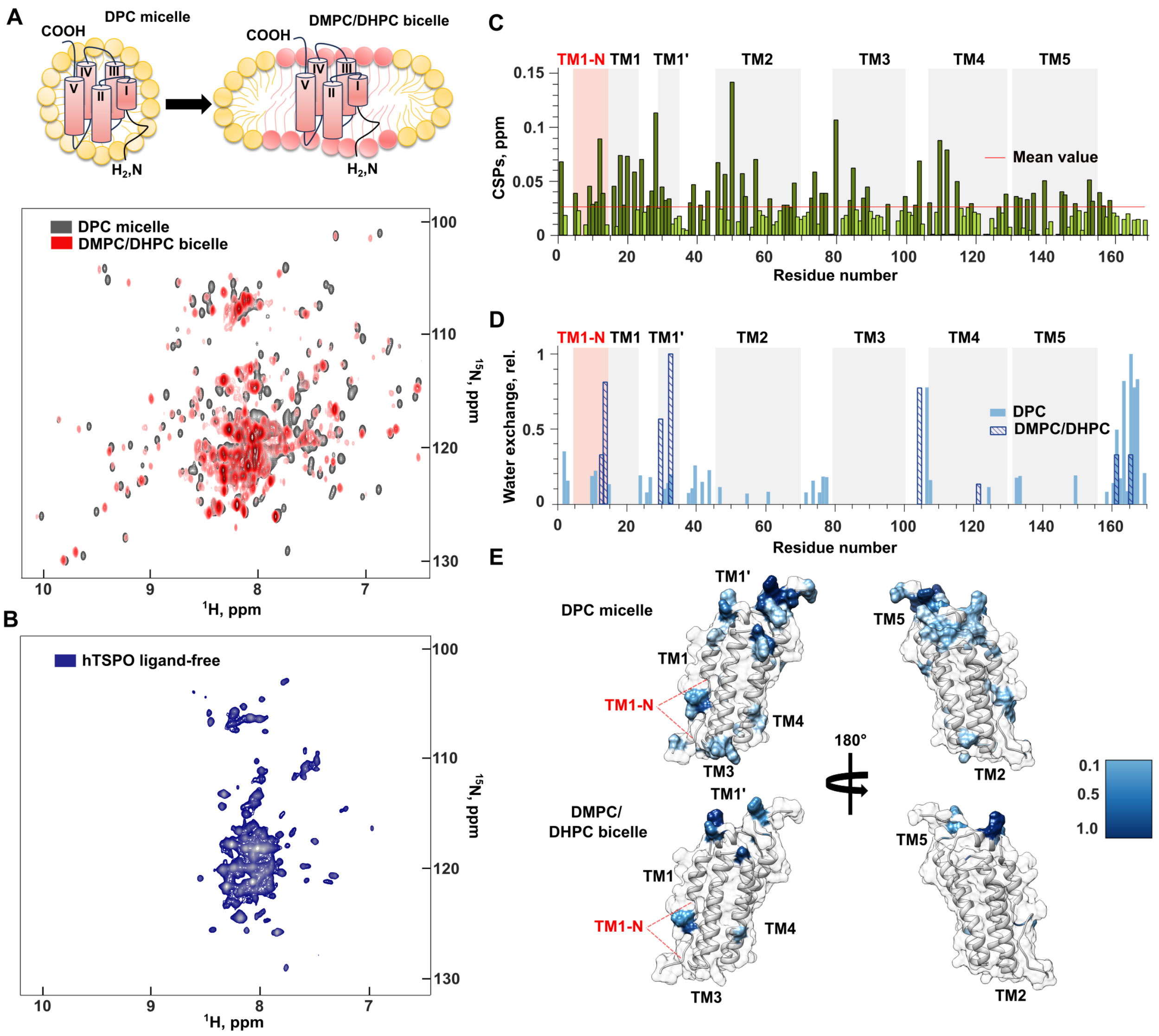


**Fig. S2. Structural properties of human TSPO in bicelles and micelles.** (**A**) Schematic representation of hTSPO reconstitution in micelles and bicelles (top). Superposition of 2D [$^{1}$H,$^{15}$N]-TROSY spectra of hTSPO/GE-180 in DPC micelles (black) and isotropic bicelles (red; DMPC/DHPC, $q = 0.5$) (bottom). (**B**) [$^{1}$H,$^{15}$N]-TROSY spectra of hTSPO in DPC micelles. (**C**) Residue-specific chemical shift perturbations (CSPs) between DPC micelles and DMPC/DHPC bicelles. The mean CSP value (red; 0.028 ppm) is indicated. Light green marks residues below the mean, and dark green those above it. (**D**) Residue-specific amide–water NOE contact profiles in micelles and bicelles, shown as bar plots. (**E**) Mapping of amide–water NOE contacts onto the hTSPO structure in micelles (top) and bicelles (bottom) reveals broadly similar patterns, with TM1-N maintaining water contacts in both environments.

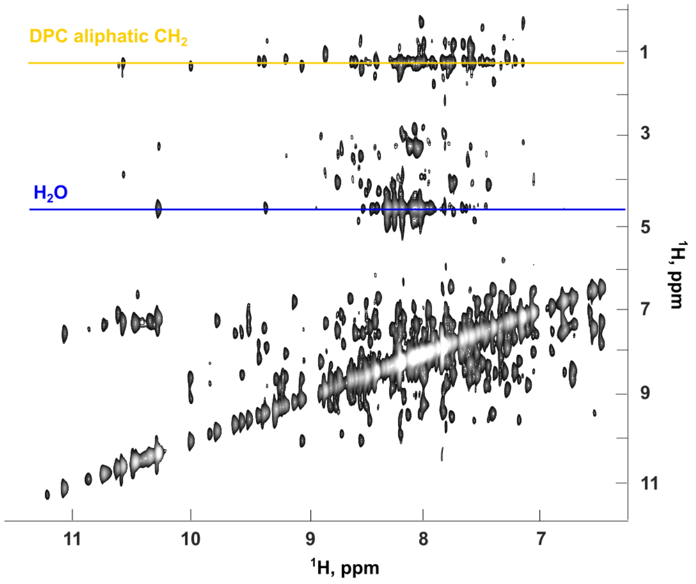


**Fig. S3. $^1H$-$^1H$ projection of the 3D $^{15}N$-edited NOESY-TROSY spectrum of hTSPO–GE-180 in DPC micelles.** The projection onto the $^1H$–$^1H$ plane highlights intermolecular NOEs between protein backbone amide protons and either water (~4.7 ppm, blue line) or the aliphatic $CH_2$ protons of DPC (1.1–1.3 ppm, yellow line). These cross-peaks were used to quantify residue-specific water and detergent contacts.

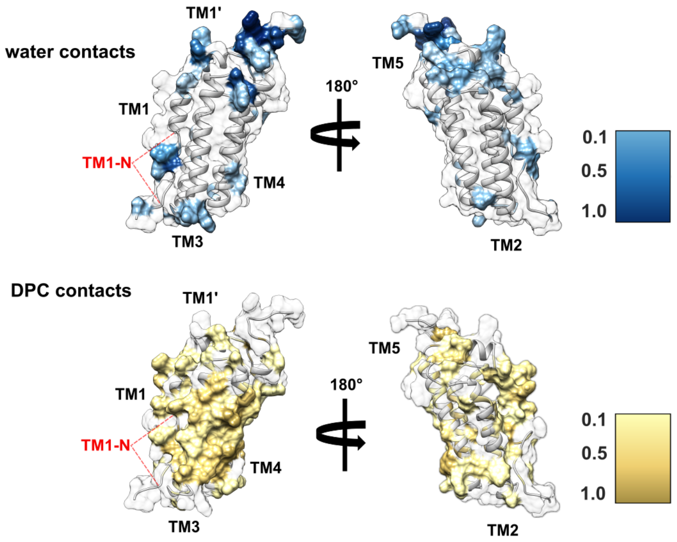


**Fig. S4. Structural mapping of water and detergent contacts in hTSPO.** Surface representation of hTSPO colored according to relative amide–water and amide–DPC NOE intensities derived from 3D $^{15}$N-edited NOESY-TROSY spectra. Loops and the N- and C-termini show strong water contacts, whereas the transmembrane domains are mainly characterized by detergent (DPC) interactions. The TM1-N segment is highly accessible to both water and detergent, consistent with its flexible nature.

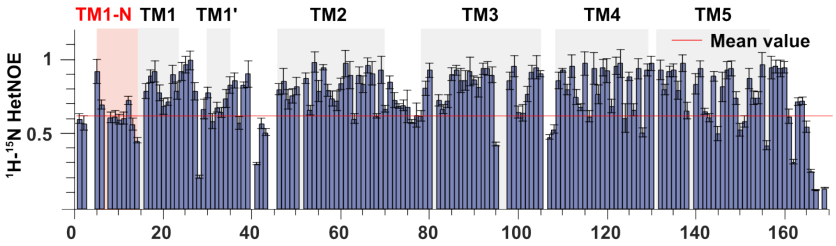


**Fig. S5.** $^{15}$N hetNOE values for hTSPO; residues with hetNOE < 0.62 (mean) were excluded from the $R_1R_2$ analysis.

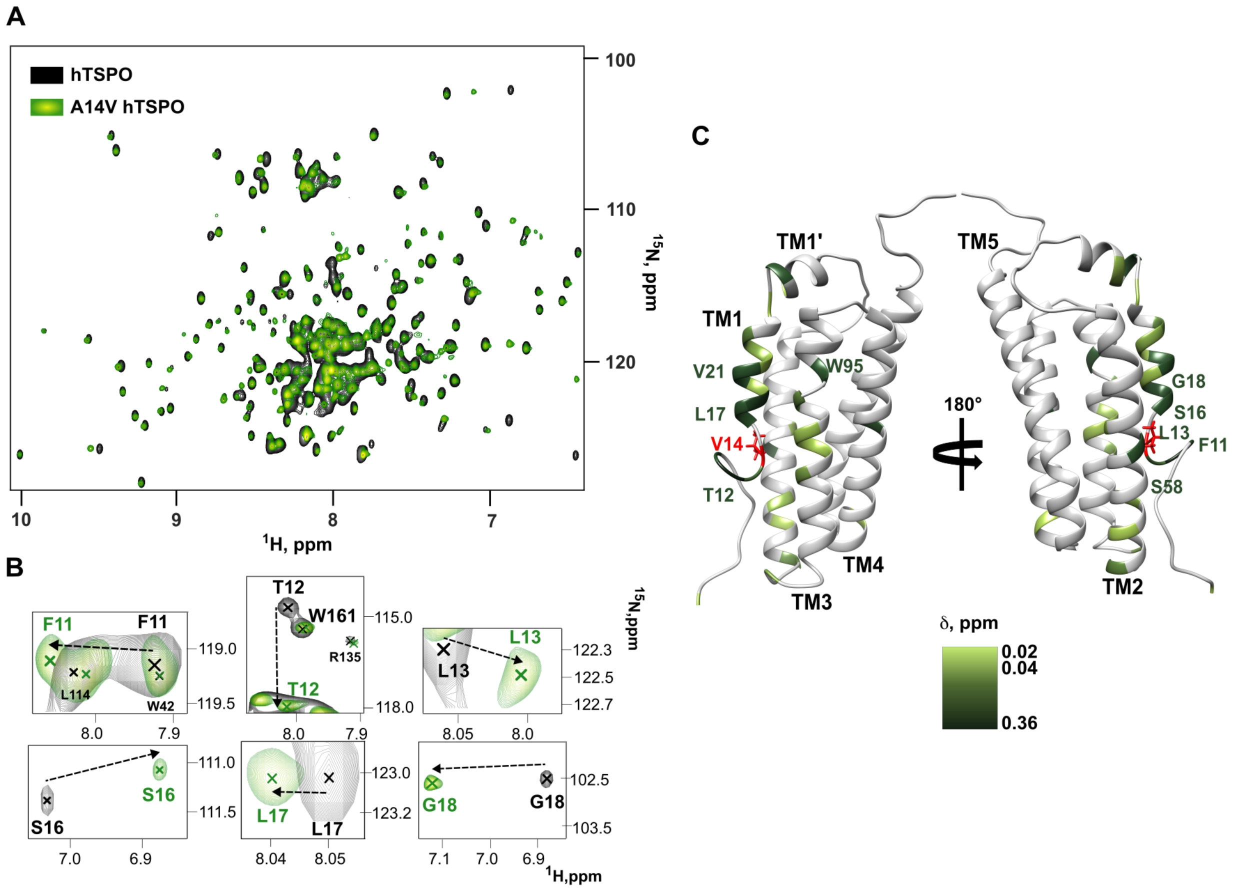


**Fig. S6. TROSY-HSQC comparison of wild-type and A14V hTSPO.** (**A**) Overlay of [$^{1}$H,$^{15}$N]-TROSY HSQC spectra of hTSPO in complex with GE-180 in DPC micelles: wild-type (black) and A14V (green). The spectra are well dispersed and overall similar. (**B**) Zoomed regions highlighting backbone amide signals with the largest CSPs near the mutation site in TM1 (examples: F11, T12, L13, S16, L17, G18). Crosses mark peak positions; dashed arrows indicate the displacement from wild-type to A14V. (**C**) CSPs between wild-type and A14V hTSPO mapped onto the protein structure.

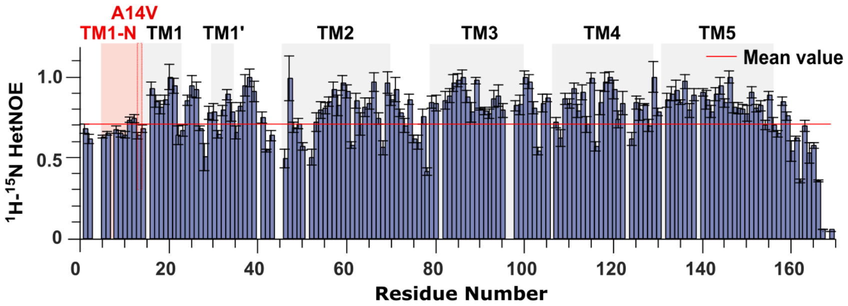


**Fig. S7.** $^{15}N\{^{1}H\}$ hetNOE values for A14V hTSPO; residues with hetNOE < 0.67 (mean) were excluded from the $R_1R_2$ analysis.

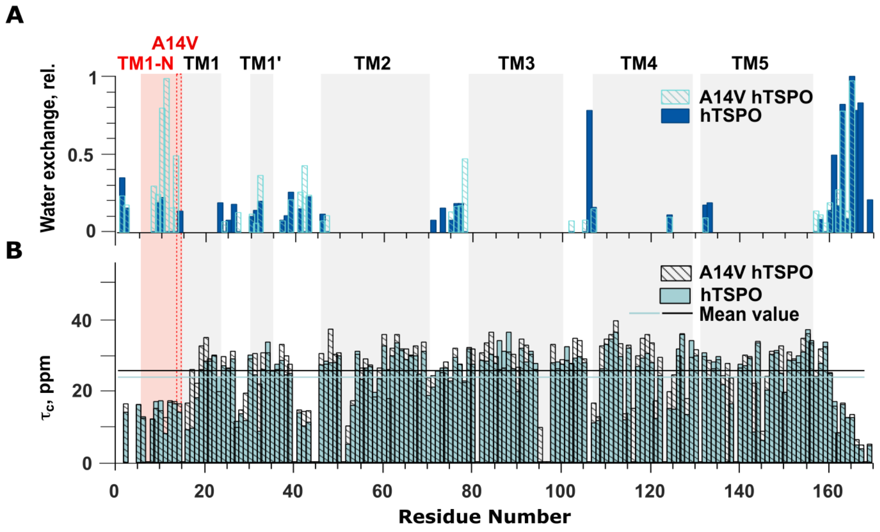


**Fig. S8.** (**A**) Residue-specific amide–water NOE profiles of wild-type hTSPO and A14V hTSPO in DPC micelles. TM1-N remains solvent-exposed in both variants. (**B**) Correlation times ($\tau_c$) for wild-type and A14V hTSPO. The average $\tau_c$ values are 24.1 ns for wild-type and 25.7 ns for A14V hTSPO. The higher average $\tau_c$ is consistent with slower overall reorientation of the protein.

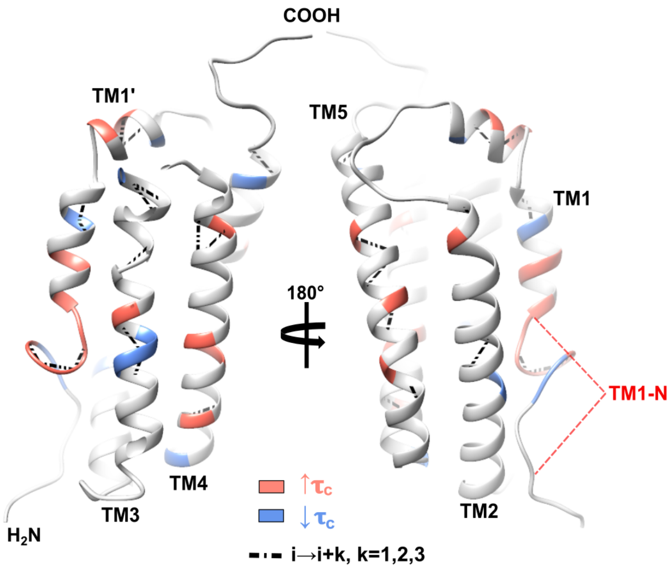


**Fig. S9. A14V induces local stabilization and short-range contacts that co-localize with changes in backbone dynamics.** Residues showing increased (red) or decreased (blue) rotational correlation times ($\Delta\tau_c$) are mapped onto the hTSPO structure. Regions exhibiting new short-range i→i+k contacts (k = 1-3) overlap with sites of altered $\tau_c$, indicating mutation-induced redistribution of backbone dynamics within the transmembrane bundle.

**Table S1. NMR experiments and acquisition parameters for WT and A14V hTSPO samples.**

| Experiment | Pulse Program | Field (MHz) | Resolution | NS | Notes |
|---|---|---|---|---|---|
| **hTSPO in DPC** | | | | | |
| **HSQC** | trosyetf3gpsi | 900 | 2048×256 | 32 | |
| **HNCA** | trhncaetgp3d | 900 | 2048×60×80 | 56 | NUS |
| **HNCO** | trhncocaetgp3d | 600 | 2048×70×84 | 64 | |
| **HNCACB** | trhncacbetgp3d | 900 | 2048×92×66 | 56 | |
| **HNcoCA** | trhncocaetgp3d | 800 | 2048×78×80 | 32 | NUS |
| **HNcaCO** | trhncacoetgp3d | 600 | 2048×70×72 | 72 | |
| **NOESY-TROSY** | noesytretf3gp3d | 1200 | 2048×64×128 | 56 | NUS, mixing time 120ms |
| **$T_1$ ($^{15}$N)** | *(39)* | 950 | 2048×128 | 42 | Delays: 0.2s, 0.4s, 0.56s, 1.0s, 1.52s, 2.2s, 3.04s |
| **$T_2$ ($^{15}$N)** | *(39)* | 950 | 2048×128 | 42 | Delays: 8ms, 16ms, 24ms, 32ms, 40ms, 48ms, 64ms, 96ms, 128ms, 160ms, 192ms, 256ms. |
| **hetNOE ($^{15}$N)** | *(39)* | 950 | 2048×128 | 42 | |
| **$^{15}$N-{$^1$H} cross-correlation** | *(40)* | 950 | 2048×208 | 256 | Constant-time delay 5.4ms |
| **hTSPO in DMPC/DHPC** | | | | | |
| **HSQC** | trosyetf3gpsi | 900 | 2048×256 | 1328 | |
| **NOESY-TROSY** | noesytretf3gp3d | 950 | 2048×54×120 | 192 | NUS, mixing time 120ms |
| **A14V hTSPO in DPC** | | | | | |
| **HSQC** | trosyetf3gpsi | 950 | 2048×256 | 32 | |
| **HNCA** | trhncaetgp3d | 950 | 2048×60×80 | 64 | NUS |
| **HNCO** | trhncocaetgp3d | 600 | 2048×70×66 | 48 | |
| **HNcaCO** | trhncacoetgp3d | 600 | 2048×66×80 | 64 | |
| **NOESY-TROSY** | noesytretf3gp3d | 950 | 2048×58×120 | 32 | NUS, mixing time 120ms |
| **$T_1$ ($^{15}$N)** | *(39)* | 950 | 2048×128 | 42 | Delays: 0.2s, 0.4s, 0.56s, 1.0s, 1.52s, 2.2s, 3.04s |
| **$T_2$ ($^{15}$N)** | *(39)* | 950 | 2048×128 | 42 | Delays: 8ms, 16ms, 24ms, 32ms, 40ms, 48ms, 64ms, 96ms, 128ms, 160ms, 192ms, 256ms. |
| **hetNOE ($^{15}$N)** | *(39)* | 950 | 2048×128 | 42 | |
| **$^{15}$N-{$^1$H} cross-correlation** | *(40)* | 950 | 2048×208 | 256 | Constant-time delay 5.4ms |

All experiments were performed at 42°C. Non-uniform sampling (NUS) was not used unless explicitly stated in the "Notes" column.

**Table S2. Software used for data acquisition, processing, structural calculation and figure preparation.**

| Software / Tool | Version | Purpose in this study |
|---|---|---|
| TopSpin (Bruker BioSpin) | 3.6.3; 4.3.0 | NMR data acquisition and processing |
| NMRFx Analyst | 11.4.130 | Multidimensional NMR data processing |
| CCPNMR Analysis | 3.2.2.131 | Peak picking, resonance assignment, and relaxation analysis |
| CYANA (including FLYA module) | - | Automated backbone assignment |
| TALOS+ | - | Dihedral angle prediction from chemical shifts |
| XPLOR-NIH | 3.7 | Structure calculation by simulated annealing |
| AlphaFold3 | 2024 release | Structural modeling |
| UCSF Chimera | 1.7, X | Structural visualization |
| SciDAVis | 2.7.1 | Data fitting, plotting and quantitative analysis |
| Microsoft Excel | 365 | Data handling and statistical calculations |
| Inkscape | 1.3.2 | Figure preparation and layout |
